# The p150 Isoform of ADAR1 Blocks Sustained RLR signaling and Apoptosis during Influenza Virus Infection

**DOI:** 10.1101/2020.05.23.111419

**Authors:** Olivia A. Vogel, Julianna Han, Chieh-Yu Liang, Santhakumar Manicassamy, Jasmine T. Perez, Balaji Manicassamy

## Abstract

Signaling through retinoic acid inducible gene I (RIG-I) like receptors (RLRs) is tightly regulated, with activation occurring upon sensing of viral nucleic acids, and suppression mediated by negative regulators. Under homeostatic conditions aberrant activation of melanoma differentiation-associated protein-5 (MDA5) is prevented through editing of endogenous dsRNA by RNA editing enzyme Adenosine Deaminase Acting on RNA (ADAR1). In addition, ADAR1 is postulated to play proviral and antiviral roles during viral infections that are dependent or independent of RNA editing activity. Here, we investigated the importance of ADAR1 isoforms in modulating influenza A virus (IAV) replication and revealed the opposing roles for ADAR1 isoforms, with the nuclear p110 isoform restricting versus the cytoplasmic p150 isoform promoting IAV replication. Importantly, we demonstrate that p150 is critical for preventing sustained RIG-I signaling, as p150 deficient cells showed increased IFN-β expression and apoptosis during IAV infection, independent of RNA editing activity. Taken together, the p150 isoform of ADAR1 is important for preventing sustained RIG-I induced IFN-β expression and apoptosis during viral infection.

## Introduction

Cell intrinsic responses against RNA viruses are primarily mediated through cytoplasmic retinoic acid inducible gene I (RIG-I) like receptors (RLRs). RLRs recognize motifs present in the viral genome, with RIG-I sensing short dsRNA with 5’ triphosphate and melanoma differentiation-associated protein 5 (MDA5) detecting long dsRNA (Pichlmair et al., 2006; Hornung et al., 2006; Schlee et al., 2009; Schmidt et al., 2009; Kato et al., 2008; Pichlmair et al., 2009; Peisley et al., 2012). Upon ligand activation, RIG-I or MDA5 translocate to the mitochondrial membrane and interact with mitochondrial antiviral signaling protein (MAVS) (Kawai et al., 2005). Tank binding kinase (TBK1) and inhibitor of nuclear factor kappa B kinase (IKKb) are then recruited to the RLR-MAVS complex to phosphorylate and activate the transcription factors interferon regulator factor 3 (IRF3) and nuclear factor kappa B (NF-kB), which subsequently translocate to the nucleus to initiate transcription of interferon-β (IFN-β) and other proinflammatory cytokines, respectively (Xu et al., 2005; Seth et al., 2005; Hou et al., 2011; Goubau et al., 2013). Secreted IFN signals in an autocrine or paracrine manner by binding to IFN receptors and activating the Janus kinase (JAK) and signal transducer and activators of transcription (STAT) pathway, which results in transcriptional upregulation of >300 interferon stimulated genes (ISGs) (Schneider et al., 2014). Through a variety of mechanisms, these induced ISGs restrict viral replication in infected cells as well as create an antiviral state in the surrounding cells.

In addition to RLR mediated transcriptional upregulation of antiviral responses, RLR activation can induce cell intrinsic apoptosis through RLR-induced IRF3 Mediated Pathway of Apoptosis (RIPA) (Chattopadhyay et al., 2017; Chattopadhyay et al., 2016; Chattopadhyay et al., 2010; Chattopadhyay et al., 2011). In this pathway, subsequent to RIG-I activation, IRF3 is activated by linear polyubiqitinylation by the linear ubiquitin chain assembly complex (LUBAC) (Chattopadhyay et al., 2016). Polyubiquitinylated IRF3 in complex with proapoptotic factor Bax translocates to the mitochondrial membrane and stimulates the release of cytochrome C, thereby initiating the cell death process to restrict viral replication.

Although viral nucleic acids are potent stimulators of RLRs, recent studies suggest that endogenous RNA encoded from the host genome can bind and stimulate RLRs. As such, RLRs are tightly regulated and are prevented from aberrant activation through negative regulation under homeostatic conditions. The RNA editing enzyme Adenosine Deaminase Acting on RNA (ADAR1) has been show to suppress MDA5 sensing of endogenous dsRNA transcribed from the *Alu* elements in the genome (Mannion et al., 2014; Pestal et al., 2015; Liddicoat et al., 2015). Under physiological conditions, ADAR1 prevents the sensing of endogenous dsRNA ligands by editing and destabilizing dsRNA structures (Liddicoat et al., 2016; Vitali et al., 2010). As such, ADAR1 deficiency results in embryonic lethality in mice, with embryos showing higher expression of ISGs and increased apoptosis of hematopoietic cells (Pestal et al., 2015; Liddicoat et al., 2015; Mannion et al., 2014; Hartner et al., 2009). In agreement with aberrant MDA5 signaling, embryonic lethality of ADAR1 KO mice is rescued by concurrent deletion of MDA5 or MAVS, with survival extending to shortly after birth (Mannion et al., 2014; Pestal et al., 2015; Liddicoat et al., 2015). Interestingly, mice expressing an editing mutant of ADAR1 are viable under MDA5 or MAVS deficient conditions, suggesting an editing dependent role of ADAR1 for survival (Liddicoat et al., 2015).

ADAR1 is expressed as two isoforms: a constitutively expressed short nuclear isoform p110 and an inducible long cytoplasmic isoform p150, which is induced by viral infection as well as by treatment with IFN (George et al., 2005; Patterson and Samuel 1995; Patterson, Thomis, et al., 1995; Shtrichman et al., 2002). Considering the RNA editing functions of ADAR1 and its role in innate immunity, there is great interest in understanding the potential proviral and antiviral roles of ADAR1 during viral infection. As several RNA viral genomes show signatures of ADAR1 editing (A→G mutations), it is postulated that regulated ADAR1 editing of viral RNA may benefit viral replication, whereas hyper editing of the viral RNA genome can reduce fitness (Cattaneo et al., 1988; Murphy et al., 1991; Martinez et al., 1997; Tenoever et al., 2007; Zahn et al., 2007; Suspene et al., 2008; Chambers et al., 2009; Taylor et al., 2005). For hepatitis delta virus, p110 editing of viral RNA is critical for production of the large antigen, signaling the switch from replication to genome packaging (Jayan et al., 2002; Wong et al., 2002). ADAR1 has been shown to be proviral for HIV, as knockdown of ADAR1 decreased LTR transcription (Phuphuakrat et al., 2008; Clerzius et al., 2009; Doria et al., 2009). For measles virus, ADAR1 editing of the genome is suggested to decrease immune stimulatory RNA (Cattaneo et al., 1988; Cattaneo et al., 1986; Pfaller et al., 2018). Additionally, knockdown or knock out of ADAR1 increased activation of PKR upon mutant measles virus infection, indicating a proviral role (Pfaller et al., 2018; Toth et al., 2009; Li et al., 2012; Okonski et al., 2013). In contrast, ADAR1 is suggested to be antiviral for HCV, as knockdown of ADAR1 increased viral RNA levels (Taylor et al., 2005). In the context of influenza A virus (IAV), ADAR1 has been proposed to play both proviral and antiviral roles; knockdown of ADAR1 in human lung epithelial cells (A549s) reduced viral replication, suggesting a proviral role for ADAR1 in influenza virus replication (de Chassey et al., 2013). In contrast, IAV showed increased cytopathic effect in p150 KO mouse embryonic fibroblasts, suggesting an antiviral role for p150 during influenza virus replication (Ward et al., 2011). Interestingly, ADAR1 has been shown to interact with the IAV non-structural protein 1 (NS1) through yeast two-hybrid screens (de Chassey et al., 2013; Ngamurulert et al., 2009). While the exact function of ADAR1 in IAV replication has yet to be fully elucidated, these studies highlight the potential for ADAR1 as a host factor involved in IAV replication.

Here, we investigated the role of ADAR1 in IAV infection by generating ADAR1 and isoform specific KOs in A549s using the CRISPR/Cas9 technology. We demonstrate that the loss of ADAR1 or p150 isoform increased expression of ISGs under homeostatic conditions, due to MDA5-mediated sensing of endogenous ligands. Although concurrent deletion of p150 and MDA5 reduced basal IFN-β expression, IAV replication was significantly reduced in p150/MDA5 double KOs (DKOs) due to sustained IFN-β expression and increased cell intrinsic apoptosis. Both elevated IFN receptor signaling via the Jak-STAT pathway and increased apoptosis contributed to restriction of IAV replication in various cells deficient in p150. The restoration of IAV replication in p150/MDA5/RIG-I and p150/MDA5/MAVS triple KOs (TKOs) demonstrate that p150 prevents sustained RIG-I activation induced IFN-β expression and apoptosis. Importantly, we demonstrate that the RNA binding activity, not the editing activity, of p150 is critical for the suppression of RIG-I signaling during IAV infection. Together, these results demonstrate that p150-mediated suppression of RIG-I signaling creates a more hospitable environment for efficient IAV replication.

## Results

### ADAR1 is a proviral factor for IAV replication

To assess the role of ADAR1 in IAV replication, we generated CRISPR/Cas9 knockouts of ADAR1 in A549s (ADAR1 KOs) using a single guide RNA (gRNA) targeting the open reading frame shared by both isoforms (Figure 1A). Individual ADAR1 KO clones were identified by Sanger sequencing of the region flanking the sgRNA target site. To confirm the loss of ADAR1 protein expression, we performed western blot analysis for the p110 and p150 isoforms in ADAR1 KOs, vector control A549s (CTRL), and parental wild-type A549s (WT A549) under mock and universal interferon (IFN-I) treatment conditions, which induces p150 expression (Figure 1B). ADAR1 KOs showed loss of expression for both p110 and p150 under mock and IFN-I treated conditions, confirming successful knockout of both isoforms. Next, to assess the importance of ADAR1 in IAV replication, ADAR1 KOs and CTRL A549s were infected at a low MOI with different strains of IAV (A/Puerto Rico/8/1934 H1N1, A/Vietnam/1203/04 H5N1-low pathogenic, and A/HongKong/1/1968 H3N2) and observed decreased IAV replication in ADAR1 KOs as compared to CTRL A549s (Figure 1C – Figure Supplement 1A). Interestingly, the replication of vesicular stomatitis virus (VSV), also an RNA virus, was similar in both ADAR1 KOs and CTRL A549s (Figure 1C). These results suggest that ADAR1 is important for optimal IAV replication.

**Figure 1.**
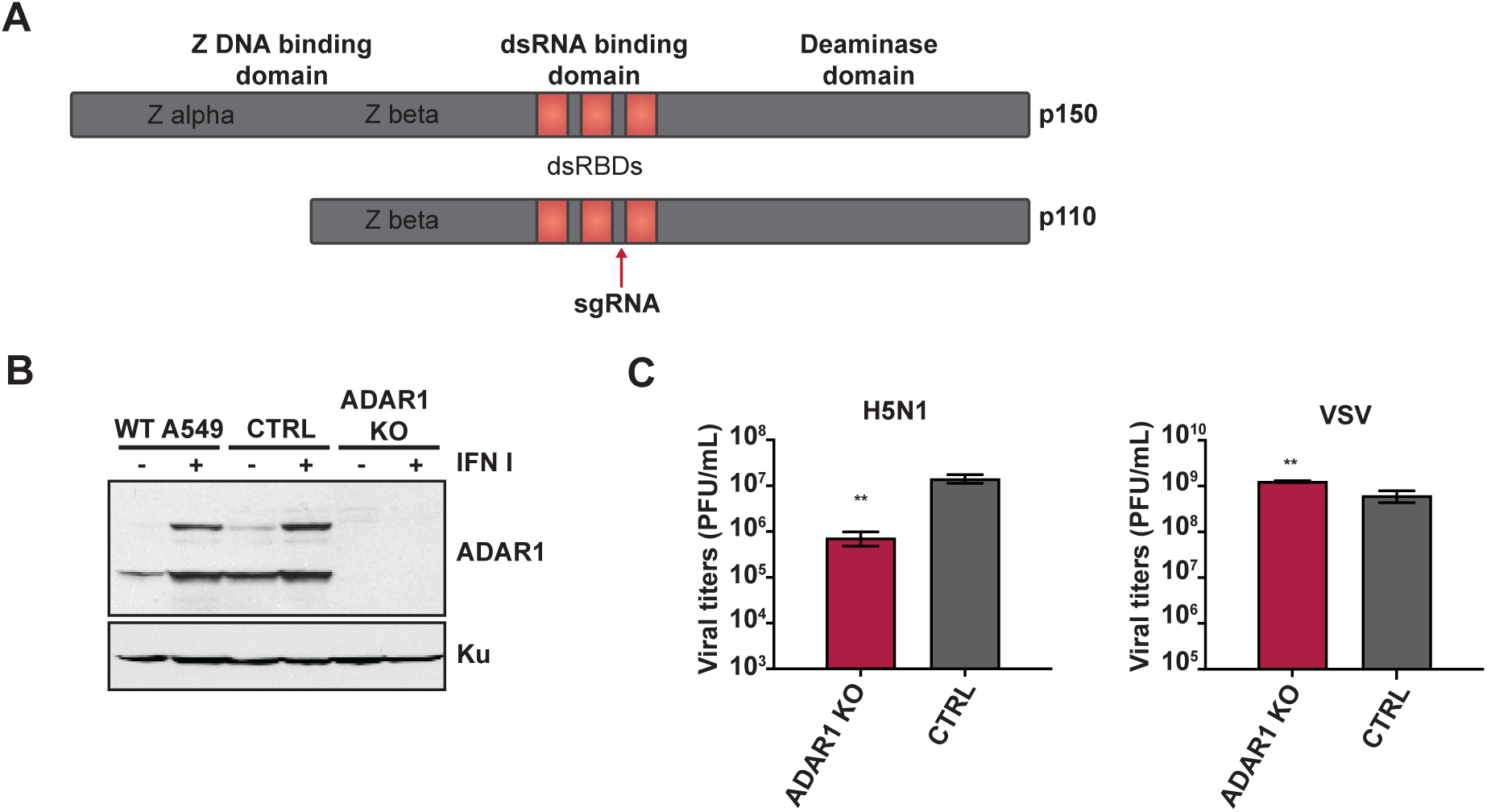
ADAR1 is an important host factor for IAV replication. (A) Schematic representation of the p110 and p150 isoforms of ADAR1, including the functional domains in each isoform. The red arrow upstream of the third dsRNA binding domain indicates the region targeted by the sgRNA in ADAR1. (B) Western blot analysis of ADAR1 expression in ADAR1 KO and control cells. ADAR1 KO, CTRL, and WT A549s were mock treated or treated with IFN for 24 hours and expression of ADAR1 was examined by western blot. Expression of Ku is shown as a loading control. (C) IAV replication in ADAR1 KOs. ADAR1 KOs and CTRL A549s were infected with H5N1 (MOI = 0.001) or VSV (MOI = 0.001) and viral titers were measured at 48 hours. Data are represented as mean titer of triplicate samples ± SD. * denotes p-value *≤* 0.5. ** denotes p-value *≤* 0.01. *** denotes p-value *≤* 0.001. NS denotes p-value *≥* 0.05. Data are representative of at least three independent experiments. See also Figure Supplement 1.

### The p150 isoform of ADAR1 is important for optimal IAV replication

Next, to identify the isoform of ADAR1 promoting IAV replication, we generated isoform specific KO A549s by deleting either the p110 or p150 promoter (p110 KOs or p150 KOs, respectively) using sgRNA (Figure 2A). Individual KO clones were identified by PCR screening for deletion and the loss of specific ADAR1 isoforms was analyzed by western blot analysis following IFN-I treatment (Figure 2B). Under mock conditions, p110 expression was abolished in p110 KOs but not in p150 KOs or CTRL A549s. Following treatment with IFN-I, p150 KOs did not show induction of p150 expression, confirming successful deletion of the p150 promoter. In the p110 KOs, treatment with IFN-I led to induction of both p110 and p150 expression, albeit at lower levels as compared to p150 KOs and CTRL A549s. This is in accordance with a prior study suggesting that p110 can be expressed from a cryptic promoter present before exon 1C (Figure 2A) (Chung et al., 2018; Nachmani et al., 2014).

**Figure 2.**
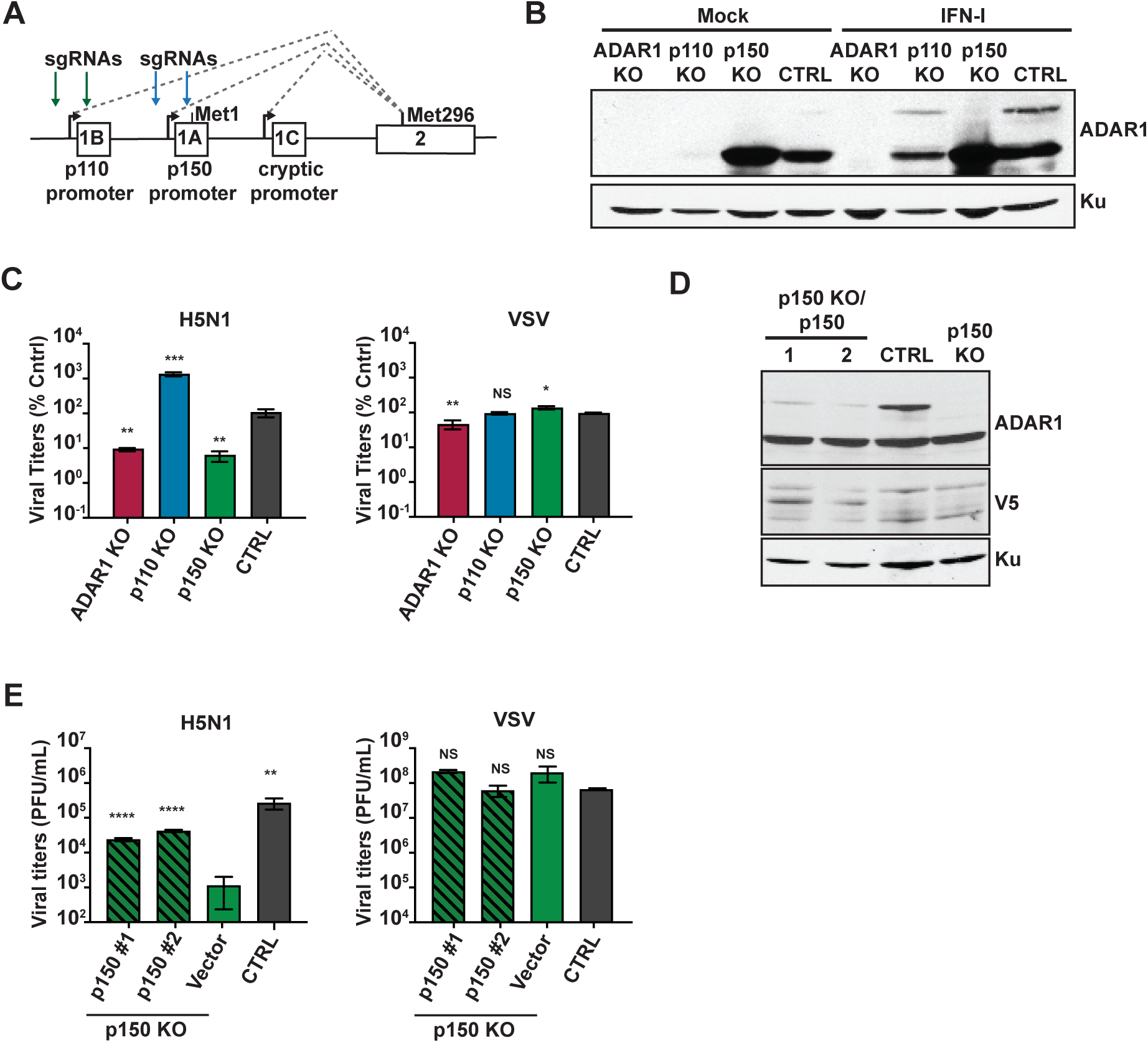
The p150 isoform of ADAR1 is critical for IAV replication. (A) Schematic representation of the promoters for the p110 and p150 isoforms of ADAR1. The promoter for p110 is upstream of exon 1B and the start methionine for p110 is within exon 2. The promoter for p150 is upstream of exon 1A and the start methionine for p150 is within exon 1A. The green arrows depict the sgRNA target sites for p110 KO and the blue arrows depict the sgRNA target sites for p150 KO. (B) Western blot analysis of ADAR1 expression in isoform specific KO A549s. ADAR1 KO, p110 KOs, p150 KOs, and CTRL A549s were mock treated or treated with IFN for 24 hours and ADAR1 expression was examined by western blot. Expression of Ku is shown as a loading control. (C) IAV replication in isoform specific KO A549s. ADAR1 KOs, p110 KOs, p150 KOs, and CTRL A549s were infected with H5N1 (MOI = 0.001) or VSV (MOI = 0.001) and viral titers were measured at 48 hours. (D) Western blot analysis of V5 tagged p150 in complemented p150 KO A549s. Two clones of p150 complements p150 KO (p150KO/p150 #1, #2), p150 KO/Vector and CTRL A549s were analyzed for expression of ADAR1 and V5 by western blot. Expression of Ku is shown as a loading control. (E) IAV replication in p150 KO/p150#1. p150 KO/p150#2, p150 KO/Vector, and CTRL A549s were infected with H5N1 (MOI = 0.001) and VSV (MOI = 0.001) and viral titers were measured at 48 hours. Data are represented as mean titer of triplicate samples ± SD. * denotes p-value *≤* 0.5. ** denotes p-value *≤* 0.01. *** denotes p-value *≤* 0.001. NS denotes p-value *≥* 0.05. Data are representative of at least three independent experiments. See also Figure Supplement 2.

Next, we assessed IAV replication in the isoform specific KOs, ADAR1 KOs, and CTRL A549s (Figure 2C – Figure Supplement 2A). The replication of all three IAV strains was decreased in p150 KOs to levels similar to ADAR1 KOs, demonstrating that the p150 isoform is critical for optimal IAV replication. In a multi-cycle growth kinetics assay, we also observed reduced viral titers in the p150 KOs as compared to CTRL A549s at all tested time points (Figure Supplement 2B). In contrast, IAV replication was increased in p110 KOs as compared to CTRL A549s, indicating a potential antiviral role for p110 during IAV infection (Figure 2C – Figure Supplement 2A). However, VSV replication was mostly unaffected by the loss of either the p110 or p150 isoform.

Next, to confirm that the reduction in IAV replication observed in p150 KOs was due to loss of p150 expression, as well as to rule out any off-target effects, we complemented the p150 KO with V5-tagged wildtype p150 (p150 KO/p150) and confirmed the expression of exogenous p150 by western blot analysis of clonal cells (Figure 2D). As anticipated, IAV replication was increased in p150 KO/p150 clones as compared to empty vector p150 KOs for three different IAV strains, indicating that the decreased IAV replication observed in p150 KOs was due to loss of p150 expression (Figure 2E – Figure Supplement 2C). Together, these results demonstrate that the two isoforms of ADAR1 have opposing roles during IAV replication, with the p150 isoform of ADAR1 being important for efficient IAV replication, and the p110 isoform playing a potential antiviral role in IAV replication.

### p150 promotes IAV replication independent of MDA5 suppression

Prior studies indicate that loss of ADAR1 results in increased upregulation of IFN-β and ISGs due to MDA5 mediated sensing of endogenous dsRNA (Liddicoat et al., 2015; Pestal et al., 2015). To test this, we first assessed basal IFN-β expression in ADAR1 and isoform specific KOs by qRT-PCR (Figure 3A). As compared to CTRL A549s, ADAR1 KOs and p150 KOs exhibited elevated levels of basal IFN-β expression, confirming the role of p150 in suppressing MDA5 activation by endogenous ligands (Pestal et al., 2015; Liddicoat et al., 2015; Mannion et al., 2014). To determine whether the elevated basal IFN-β expression observed in p150 KOs contributed to reduced IAV replication, we generated p150 and MDA5 double knockout A549s as well as MDA5 single KO A549s (p150/MDA5 DKOs or MDA5 KOs). We confirmed successful knockout of MDA5 expression by western blot analysis (Figure 3B). As expected, p150/MDA5 DKOs exhibited reduced basal IFN-β expression as compared to p150 KOs (Figure 3C), further confirming that p150 suppresses sensing of endogenous ligands by MDA5. Next, to determine if the reduced basal IFN-β levels rendered p150/MDA5 DKOs more permissive to IAV replication, we assessed IAV replication in various MDA5 KOs. Interestingly, IAV replication was reduced in p150/MDA5 DKOs to levels similar to p150 KOs (Figure 3D – Figure Supplement 3A). We observed a similar reduction in IAV replication in p150/MDA5 DKOs in multicycle replication assays for both H1N1 and H5N1 strains (Figure Supplement 3B). Complementation of p150/MDA5 DKOs with V5-p150 restored IAV replication to levels similar to CTRL A549s, demonstrating that the reduction in viral titer in p150/MDA5 DKOs was due to loss of p150 expression (Figure 3E). Taken together, these results demonstrate that p150 promotes IAV replication independent of MDA5 suppression.

**Figure 3.**
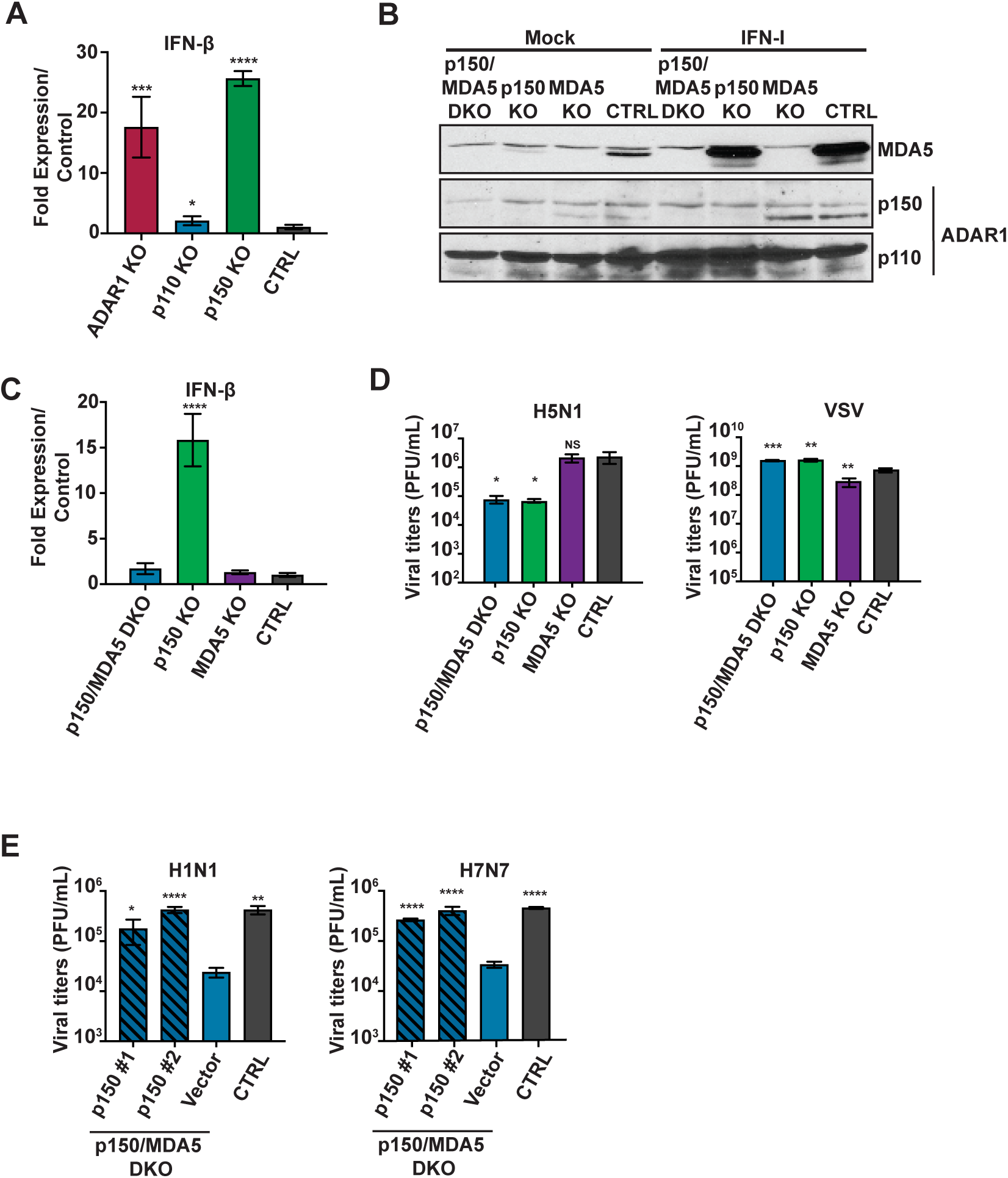
p150 supports IAV replication independent of MDA5 suppression. (A) qRT-PCR analysis of basal IFN-β expression in ADAR1 KOs, isoform specific KOs, and CTRL A549s. Data are represented as fold expression relative to vector control A549s. (B) Western blot analysis of MDA5 and ADAR1 isoforms in various KOs. p150/MDA5 DKOs, p150 KOs, MDA5 KOs, and CTRL A549s were mock treated or treated with IFN for 24 hours. ADAR1 and MDA5 expression were examined by western blot. (C) qRT-PCR analysis of basal IFN-β expression in p150/MDA5 DKOs. Data are represented as fold expression relative to CTRL A549s. (D) IAV replication in p150/MDA5 DKOs. p150/MDA5 DKOs, p150 KOs, MDA5 KOs, and CTRL A549s were infected with H5N1 (MOI = 0.001) and VSV (MOI = 0.001) and viral titers were measured at 48 hours. Data are represented as mean titer of triplicate samples ± SD. (E) IAV replication in p150/MDA5 DKOs expressing wild-type V5-tagged p150. Two clones of p150/MDA5 DKOs complemented with wildtype p150 were infected with H1N1 (MOI = 0.01) and H7N7 (MOI = 0.01). p150/MDA5 DKOs that have been transduced with empty vector and CTRL A549s were also infected. Viral titers were measured at 72 hours. Data are represented as mean titer of triplicate samples ± SD. * denotes p-value *≤* 0.5. ** denotes p-value *≤* 0.01. *** denotes p-value *≤* 0.001. NS denotes p-value *≥* 0.05. Data are representative of at least three independent experiments. See also Figure Supplement 3.

### p150 suppresses exogenous RLR ligand induced IFN-β expression and apoptosis

Prior studies indicate that RLR-mediated sensing of viral PAMPS activates both IRF3 mediated transcriptional upregulation of antiviral genes as well as apoptosis through the RIPA pathway as means to control viral replication (Figure 4A) (Chattopadhyay et al., 2010; Chattopadhyay et al., 2011; Chattopadhyay et al., 2016; Chattopadhyay et al., 2017). As we observed decreased IAV replication in p150/MDA5 DKOs, we next investigated the role of p150 in RIG-I mediated induction of IFN-β expression and apoptosis. We first examined IFN-β expression in p150/MDA5 DKOs following transfection of low molecular weight (LMW) pI:C and high molecular weight (HMW) pI:C, which predominantly stimulate RLRs RIG-I and MDA5, respectively (Figure 4B). Transfection of LMW pI:C showed elevated IFN-β expression in p150/MDA5 DKO as compared to CTRL cells; however, transfection of HMW pI:C did not result in higher IFN-β expression in p150/MDA5 DKOs due to the lack of MDA5. These results suggest that, in addition to suppressing MDA5 sensing of endogenous ligands, p150 also suppresses RIG-I mediated induction of IFN-β expression from exogenous stimuli. Next, we examined the kinetics of IFN-β expression during SeV infection, a potent activator of RLRs, in various A549 KO cells (Figure 4C). At 8 hpi, IFN-β expression levels were similar in all cell types analyzed; however, starting at 16 hpi, IFN-β expression remained elevated in p150/MDA5 DKOs and p150 KOs; in contrast, MDA5 KOs and CTRL A549s showed down regulation of IFN-β expression. To rule out altered viral replication contributing to the observed differences in IFN-β expression, we performed IAV vRNA transfections and observed sustained IFN-β expression in p150/MDA5 DKOs as compared to CTRL A549s (Figure 4D). For further validation, we generated human embryonic kidney cells lacking p150 (p150 KO 293) and examined IFN-β expression following Sendai virus (SeV) (Figure 4E – Figure Supplement 4A). IFN-β expression in the p150 KO 293s was elevated at each time point, further confirming the importance of p150 in suppressing sustained RIG-I mediated induction of IFN-β expression. Taken together, these results reveal a novel role for p150 in suppressing sustained RIG-I activation during viral infection.

**Figure 4.**
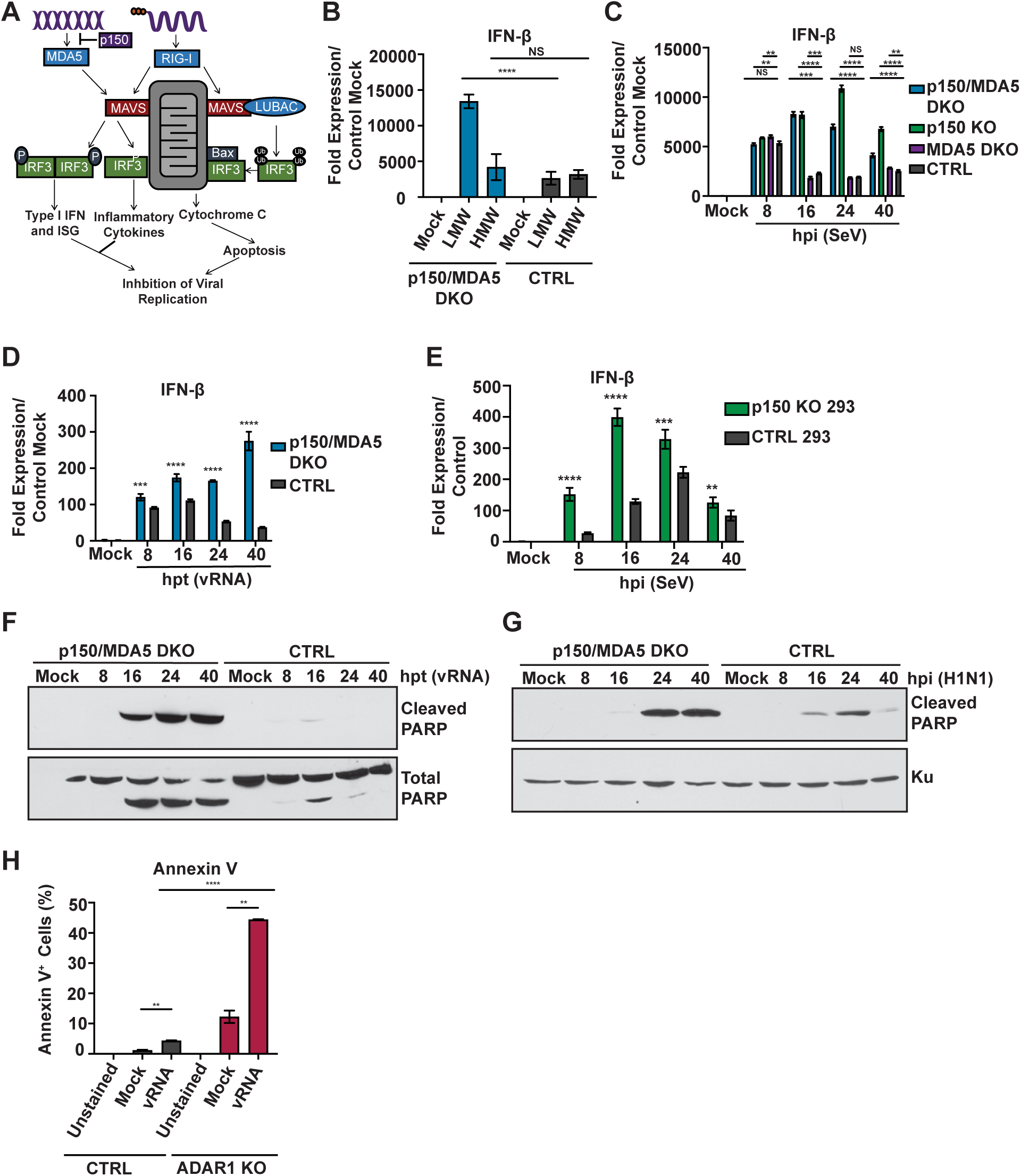
p150/MDA5 DKO A549s show elevated IFN-β expression and increased apoptosis upon RLR stimulation. (A) Schematic representation of the RLR-Induced IRF3-mediated Pathway of Apoptosis (RIPA) and IRF3 mediated transcriptional upregulation of antiviral genes. (B) qRT-PCR analysis of IFN-β expression in p150/MDA5 DKOs after poly I:C stimulation. p150/MDA5 DKOs and CTRL A549s were transfected with LWM and HMW pI:C for 24 hours and IFN-b expression was analyzed by qRT-PCR analysis. Data are represented as fold expression relative to mock transfected CTRL A549s. (C) qRT-PCR analysis of IFN-β expression in p150/MDA5 DKOs following Sendai virus (SeV). p150/MDA5 DKOs, p150 KOs, MDA5 KOs, and CTRL A549s were infected with SeV and mRNA levels were measured at the indicated hours post infection. Data are represented as fold expression relative to mock vector control. (D) qRT-PCR analysis of IFN-β expression in p150/MDA5 DKOs following H1N1 vRNA transfection. p150/MDA5 DKOs and CTRL A549s were transfected with H1N1 vRNA and mRNA levels were measured at the indicated hours post transfection (hpt). (E) qRT-PCR analysis of IFN-β expression in p150 KO and vector control 293s. p150 KO and vector control 293s were infected with SeV and mRNA levels were measured at the indicated hours post infection (hpi). Data are represented as fold expression relative to mock vector control 293s. (F-G) Western blot analysis of PARP cleavage in p150/MDA5 DKOs. (F) p150/MDA5 DKOs and CTRL A549s were transfected with H1N1 vRNA and cell lysates were collected at the indicated time points post transfection. (G) p150/MDA5 DKOs and CTRL A549s were infected with H1N1 (MOI = 1) and cell lysates were collected at the indicated time points post infection. (H) Flow cytometric analysis of Annexin V^+^ cells following H1N1 vRNA transfection. ADAR1 KOs and CTRL A549s were transfected with H1N1 vRNA and stained with Annexin V-PE 40 hours post transfection. The levels of Annexin V were analyzed by flow cytometry. * denotes p-value *≤* 0.5. ** denotes p-value *≤* 0.01. *** denotes p-value *≤* 0.001. NS denotes p-value *≥* 0.05. Data are representative of at least three independent experiments. See also Figure Supplement 4.

To assess the contribution of apoptosis via the RIPA pathway in p150 deficient cells, we examined the levels of PARP cleavage, a byproduct of apoptosis, by western blot in p150 KOs and CTRL A549s cells following stimulation with various RLR agonists (Figure Supplement 4B-D). p150 KOs exhibited increased PARP cleavage in response to IAV vRNA transfection, H1N1 infection and pI:C transfection; in contrast, little or no PARP cleavage was observed in CTRL A549s (Figure Supplement 4B-D). Similarly, p150 KO 293s exhibited increased PARP cleavage following IAV vRNA transfection (Figure Supplement 4E). These data indicate that loss of p150 results increased apoptosis upon stimulation with RLR agonists.

Next, we examined PARP cleavage in p150/MDA5 DKOs following IAV vRNA transfection or H1N1 infection (Figure 4F-G). p150/MDA5 DKOs also showed increased PARP cleavage as compared to CTRL A549s in response to both vRNA transfection and H1N1 infection. We also examined apoptosis by staining with Annexin V, which binds phosphatidylserine, an early marker for apoptosis (Figure 4H). Flow cytometric analysis of ADAR1 KOs transfected with IAV vRNA showed increased binding of Annexin V as compared to CTRL A549s. Taken together, these results demonstrate that p150 is critical for suppressing RLR induced apoptosis.

### p150 suppresses induction of IFN-β and apoptosis via the RIG-I-MAVS-IRF3 pathway

To investigate if the RIG-I signaling cascade through IRF3 results in increased apoptosis in p150/MDA5 DKOs, we performed siRNA knockdown of RIG-I or IRF3 and assessed PARP cleavage after IAV vRNA transfection (Figure 5A). Knockdown of RIG-I or IRF3 in p150/MDA5 DKOs led to reduced PARP cleavage as compared to control siRNA transfected p150/MDA5 DKOs, demonstrating that the RIG-I signaling cascade through IRF3 leads to increased apoptosis in p150 deficient cells. To further validate our findings, we generated p150/MDA5/RIG-I and p150/MDA5/MAVS triple knockout (TKO) A549s and assessed IFN-β expression and apoptosis. Following SeV infection, both the p150/MDA5/RIG-I and p150/MDA5/MAVS TKOs showed reduced IFN-β expression as compared to p150/MDA5 DKOs (Figure 5B). Similarly, both TKOs showed reduced PARP cleavage during IAV infection as compared to p150/MDA5 DKOs (Figure 5C). These data demonstrate that p150 is critical for suppressing RIG-I induced IFN-β expression and apoptosis. As expected, IAV replication in both p150/MDA5/RIG-I and p150/MDA5/MAVS TKOs increased to levels similar CTRL A549s, while replication remained lower in p150/MDA5 DKOs (Figure 5D). Taken together, these results demonstrate that the p150 isoform of ADAR1 promotes IAV replication through suppression of the RIG-I signaling cascade via MAVS-IRF3.

**Fig 5.**
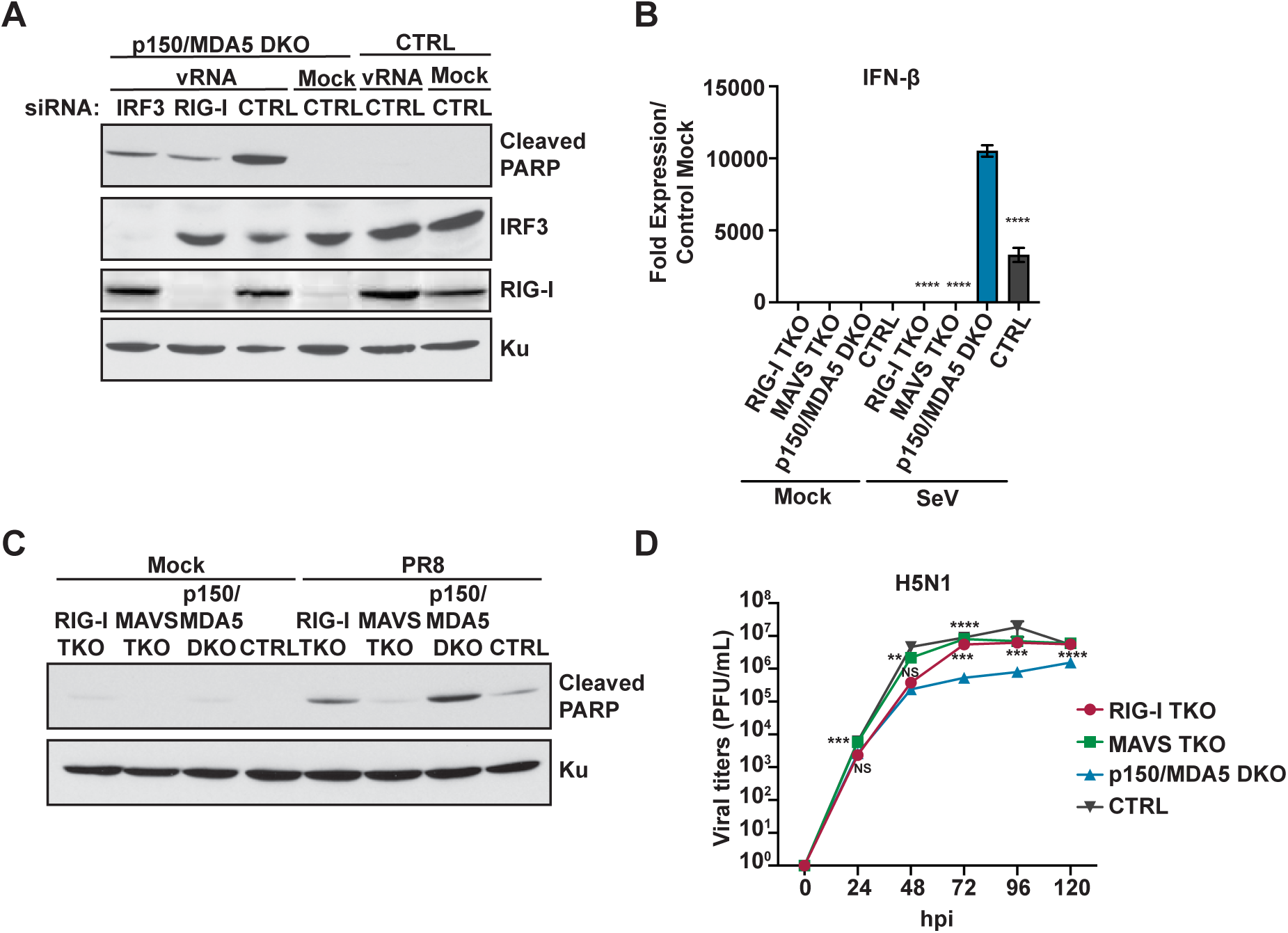
p150 suppresses RIG-I-signaling mediated induction of IFN-β and apoptosis. (A) Western blot analysis of PARP cleavage in p150/MDA5 DKOs following knockdown of RIG-I or IRF3 expression. RIG-I and IRF3 were knocked down in p150/MDA5 DKOs via siRNA transfection. Control siRNA were also transfected into p150/MDA5 DKOs and CTRL A549s. After 24h post siRNA transfection, cells were transfected with H1N1 vRNA for additional 24 hours. The cell lysates were analyzed for expression of RIG-I, IRF3, cleaved PARP, and Ku by western blot. (B) qRT-PCR analysis of IFN-β expression in p150/MDA5/RIG-I and p150/MDA5/MAVS TKOs after SeV infection. p150/MDA5/RIG-I TKOs, p150/MDA5/MAVS TKOs, p150/MDA5 DKOs, and CTRL A549s were infected with SeV and IFN-β mRNA levels were measured after 24 hours. Data are represented as fold expression relative to mock vector control A549s. (C) Western blot analysis of PARP cleavage TKOs following H1N1 infection. p150/MDA5/RIG-I TKOs, p150/MDA5/MAVS TKOs, p150/MDA5 DKOs, and CTRL A549s were infected with H1N1 (MO1 = 1) and lysates were collected after 40 hours. The levels of cleaved PARP and Ku were determined by western blot. (D) IAV replication in TKOs. p150/MDA5/RIG-I TKOs, p150/MDA5/MAVS TKOs, p150/MDA5 DKOs, and CTRL A549s were infected with H5N1 (MOI = 0.001) and viral titers were measured at the indicated time points post infection. Data are represented as mean titer of triplicate samples ± SD. * denotes p-value *≤* 0.5. ** denotes p-value *≤* 0.01. *** denotes p-value *≤* 0.001. NS denotes p-value *≥* 0.05. Data are representative of at least three independent experiments.

### Inhibition of apoptosis and Jak-STAT signaling restores IAV replication in p150 deficient cells

To determine the relative contribution of elevated IFN-β expression versus increased apoptosis to IAV restriction in p150 deficient cells, we generated p150/Bax DKO A549s using the CRISPR/Cas9 technology. Bax is a pro-apoptotic protein in the Bcl-2 protein family and has also been implicated in the RIPA pathway (Figure 4A) (Chattopadhyay et al., 2016; Chattopadhyay et al., 2017). We first assessed apoptosis in p150/Bax DKOs by PARP cleavage following IAV vRNA transfection and observed reduced PARP cleavage as compared to p150 KOs (Figure 6A). As anticipated, while PARP cleavage was reduced, IFN-β expression remained elevated in p150/Bax DKOs as compared to CTRL A549s (Figure 6B). IAV replication was increased in the p150/Bax DKOs as compared to p150 KOs yet remained lower than CTRL A549s (Figure 6C). These results indicate that inhibition of apoptosis via Bax deletion partially restores IAV replication in p150 deficient cells. Next, to assess the contribution of elevated IFN-β signaling through the IFN receptor to reduced IAV replication in p150 deficient cells, we treated various A549 KO cells with Ruxolitinib, an inhibitor of Janus kinases 1 and 2 (JAK 1/2), to block IFN receptor signaling. Treatment of p150 KOs with Ruxolitinib showed a modest increase in IAV viral titers, while Ruxolitinib treatment had no significant impact on viral replication in CTRL A549s (Figure 6D). In contrast, treatment of p150/Bax DKOs with Ruxolitinib led to a greater increase in viral titers as compared to DMSO treated p150/Bax DKOs (Figure 6D). Taken together, these results demonstrate that both increased apoptosis and elevated IFN receptor signaling contribute to the inhibition of IAV replication in the absence of p150.

**Fig 6.**
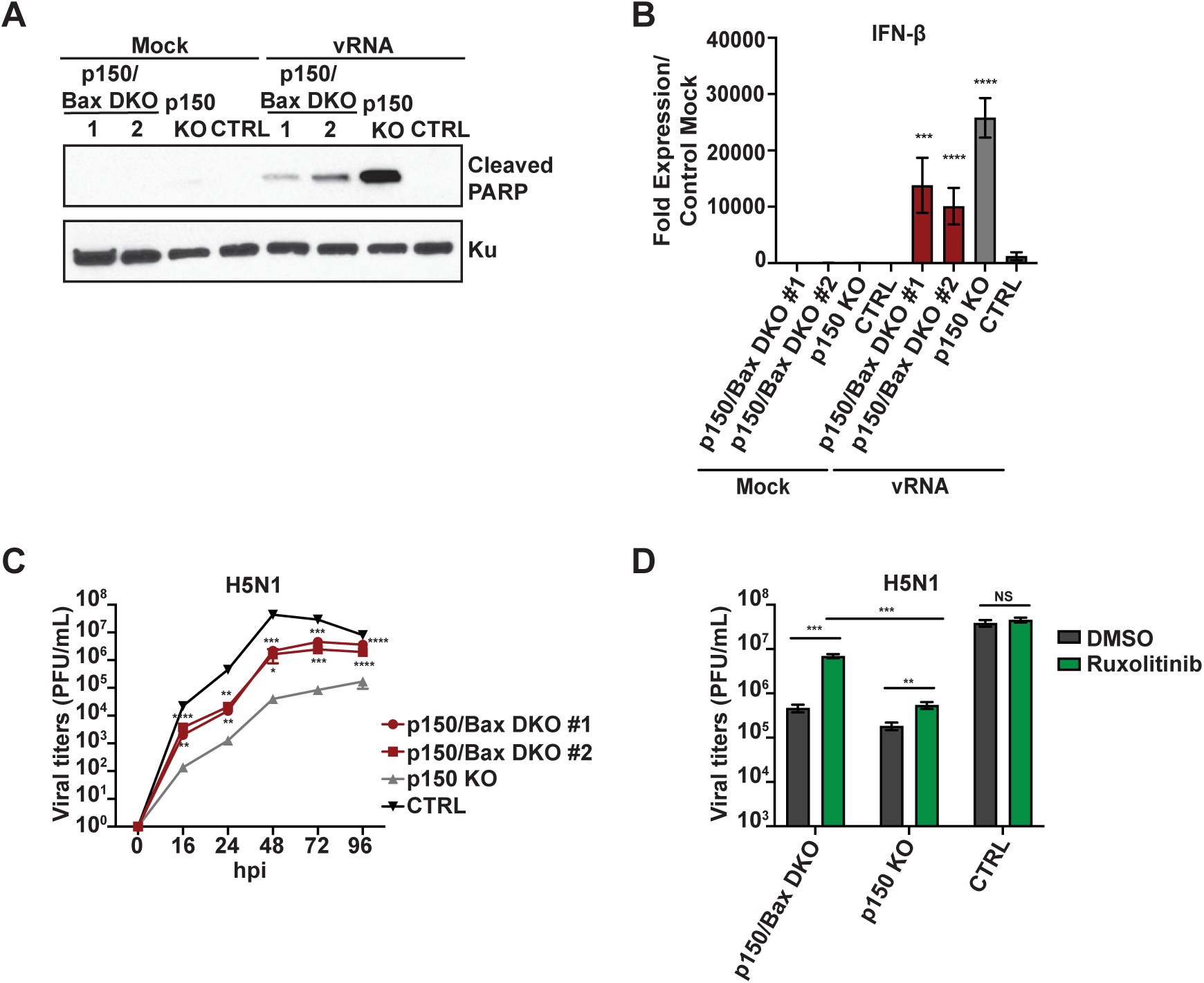
Increased type I IFN signaling and apoptosis reduce IAV replication in cells lacking p150. (A) Western blot analysis of PARP Cleavage in p150/Bax DKOs following vRNA transfection. Two clones of p150/Bax DKOs, p150 KOs, and CTRL A549s were transfected with IAV vRNA for 24 hours. PARP cleavage and Ku expression were analyzed by western blot. (B) qRT-PCR analysis of IFN-β expression in p150/Bax DKOs following IAV vRNA transfection. Two clones of p150/Bax DKOs, p150 KOs, and CTRL A549s were transfected with H1N1 vRNA and IFN-β mRNA levels were determined 24 hours post transfection. Data are represented as fold expression relative to mock vector control. (C) IAV replication in p150/BAX DKOs. Two clones of p150/Bax DKOs, p150 KOs, and CTRL A549s were infected with H5N1 (MOI = 0.001) and viral titers were measured at the indicated time points post infection. Data are represented as mean titer of triplicate samples ± SD. (D) IAV replication in p150/Bax DKOs following inhibition of Janus kinases. p150/Bax DKOs, p150 KOs, and CTRL A549s were treated with 0.2 μM Ruxolitinib for 24 hours prior to infection. The following day cells were infected with H5N1(MOI = 0.001) in the presence of Ruxolitinib and viral titers were measured at 48 hours. Data are represented as mean titer of triplicate samples ± SD. * denotes p-value *≤* 0.5. ** denotes p-value *≤* 0.01. *** denotes p-value *≤* 0.001. NS denotes p-value *≥* 0.05. Data are representative of at least three independent experiments.

### RNA binding activity of p150 is required for suppression of the RIG-I pathway

To identify the functional domains in p150 important for suppression of RIG-I signaling, we generated V5-tagged p150 mutant constructs including (1) Z alpha mutant (p150 Z*α*) with K169A/Y177A mutations in Z DNA/RNA binding domain, (2) RNA binding mutant (p150 RBM) with KKxxK→EAxxA mutations in all three RNA binding domains, and (3) a catalytic mutant that lacks the deaminase domain (p150 Cat) (Figure 7A) (Valente et al., 2007) (Ng et al., 2013). We first assessed the ability of these mutants to suppress RIG-I signaling in IFN-β reporter assay in p150 KO 293s. Different p150 mutant constructs were co-transfected with the IFN-β-firefly reporter and the RIG-I pathway was stimulated with SeV. As compared to GFP control, wildtype (WT) p150, p150 Z*α*, and p150 Cat transfected cells showed decreased IFN-β reporter activity (Figure 7B); in contrast, p150 RBM transfected cells showed increased IFN-β reporter activity. Next, we complemented the p150/MDA5 DKO A549s with different p150 mutant constructs (DKO/WT, DKO/ Z*α*, DKO/RBM, and DKO/Cat) and generated clonal populations (Figure 7C). Following IAV vRNA transfection in complemented DKOs, IFN-β expression was elevated in DKO/RBMs at levels similar to p150/MDA5 DKOs, demonstrating that the RNA binding activity of p150 is required for suppression of IFN-β expression (Figure 7D). Similarly, DKO/RBMs showed increased PARP cleavage following IAV vRNA transfection, suggesting that the RNA binding activity of p150 is also required for suppression of apoptosis (Figure 7E). Finally, we assessed IAV replication in complemented DKOs and observed increased viral titers in DKO/WTs and DKO/Cats yet not in DKO/RBMs (Figure 7F). Interestingly, DKO/Z*α*s showed decreased replication despite showing decreased IFN-β expression and apoptosis upon vRNA transfection. Taken together, these results demonstrate that the RNA binding activity and not the catalytic activity of p150 is required for the suppression of RIG-I mediated induction of IFN-β expression and apoptosis.

**Fig 7.**
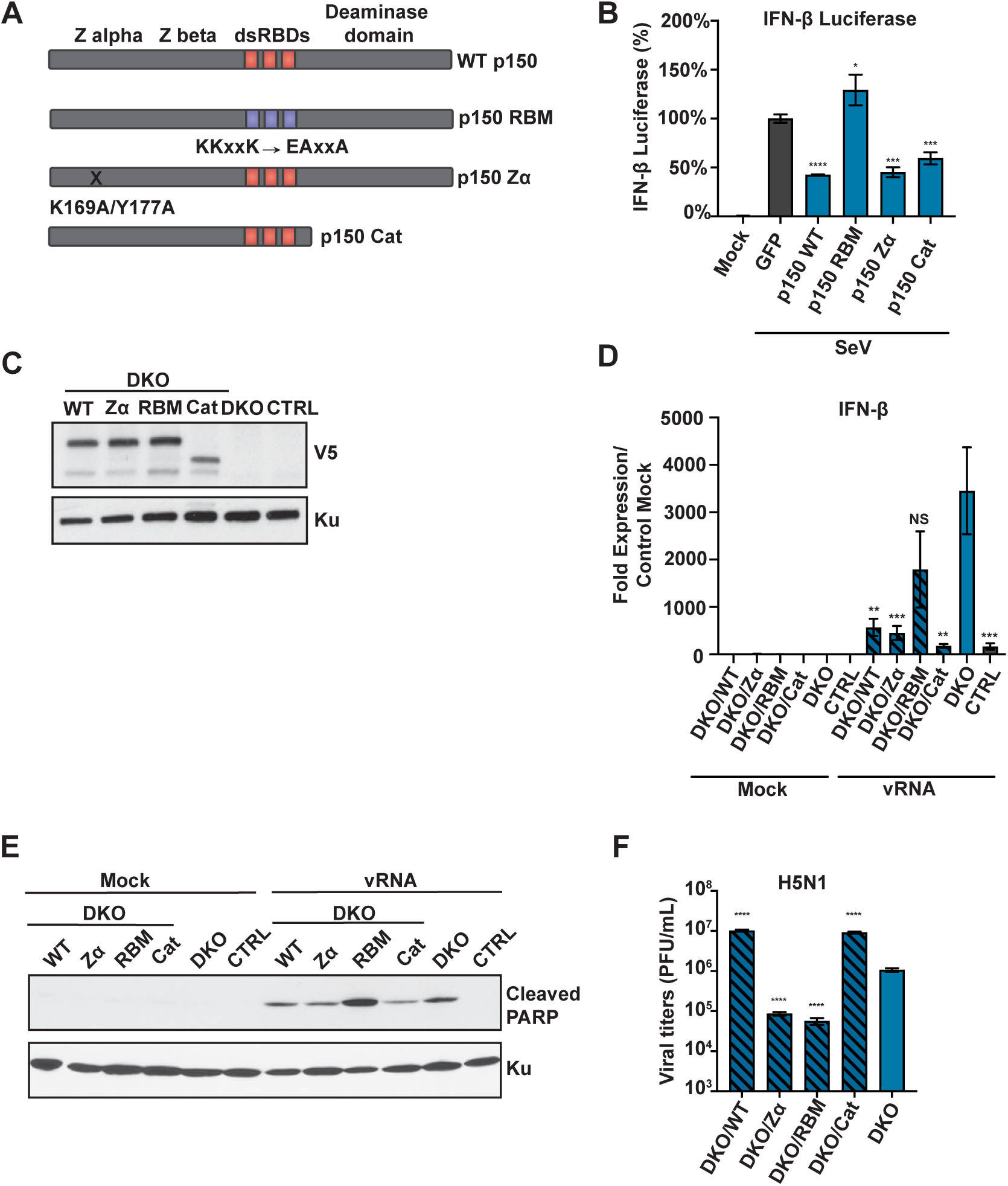
RNA binding activity of p150 is required for suppression of RIG-I signaling. (A) A schematic representation of different p150 mutants. (B) Analysis of IFN-β-Firefly reporter activity. A firefly luciferase reporter under the control of the IFN-β promoter was transfected along with wild-type and mutant p150 constructs into 293s. At 24h post-transfection, cells were infected with SeV and luciferase activity was measured 48 hours post-transfection. Data are represented as percent luciferase activity relative to GFP+SeV. (C) Western blot analysis of V5-tagged p150 expression in complemented cells. Lysates from p150/MDA5 DKOs stably expressing wild-type and mutant p150 constructs were analyzed by western blot using an anti-V5 antibody. (D) qRT-PCR analysis of IFN-β in V5-p150 complemented p150/MDA5 DKOs. p150/MDA5 DKOs complemented with different V5-p150 mutants were transfected with H1N1 vRNA and IFN-β mRNA levels were measured 24 hours post transfection. Data are represented as fold expression relative to mock CTRL A549s. (E) Western blot analysis of PARP cleavage in complemented p150/MDA5 DKOs. p150/MDA5 DKOs complemented with p150 mutants were infected with H1N1 (MOI = 1) and cell lysates were collected at 40 hours post infection for western blot analysis of cleaved PARP. (F) IAV replication in complemented p150/MDA5 DKOs. Complemented p150/MDA5 DKOs were infected with H5N1 (MOI = 0.001) and viral titers were measured at 48 hours. Data are represented as mean titer of triplicate samples ± SD. * denotes p-value *≤* 0.5. ** denotes p-value *≤* 0.01. *** denotes p-value *≤* 0.001. NS denotes p-value *≥* 0.05. Data are representative of at least three independent experiments.

## Discussion

Here, we investigated the role of ADAR1 in IAV replication and revealed opposing functions for the two isoforms, with the cytoplasmic p150 isoform acting as a negative regulator of RIG-I signaling and the nuclear p110 isoform acting as an IAV restriction factor. We show that p150 promotes IAV replication independent of its role in the suppression of MDA5 mediated sensing of endogenous dsRNA. Through concurrent deletion of p150 and RLR pathway components, we demonstrate that p150 suppresses sustained activation of the RIG-I signaling cascade, resulting in decreased levels of IFN-β expression and apoptosis. In p150 deficient cells, IAV replication was restored by inhibiting both apoptosis and IFN receptor signaling. p150 mediated suppression of RLR signaling was dependent on RNA binding functions but independent of RNA editing activity. Taken together, our studies illustrate a broad role for p150 in preventing the hyperactivation of innate immune responses, with p150 suppressing both the MDA5 pathway under basal conditions and the RIG-I pathway during viral infection.

ADAR1, a member of the ADAR family, has been implicated in several cellular functions including editing of dsRNA from repetitive elements, miRNA biogenesis/processing and regulation of cell intrinsic immunity (Athanasiadis et al., 2004; Kim et al., 2004; Blow et al., 2004; Levanon et al., 2004; Ramaswami et al., 2012; Bazak et al., 2014; Yang et al., 2006; Kawahara et al., 2008; Zipeto et al., 2016; Lamers et al., 2019). Moreover, ADAR1 has been shown to have both proviral and antiviral roles during viral infection (Samuel 2011). One well characterized proviral function of ADAR1 is in the hepatitis delta virus life cycle, where ADAR1 editing of viral RNA is critical for production of the large delta antigen (Jayan et al., 2002; Wong et al., 2002). Similarly, ADAR1 editing is critical for polyomavirus replication as it facilitates the switch from early to late gene expression (Gu et al., 2009; Kumar et al., 1997). As these viruses replicate in the nucleus, ADAR1 mediated editing of viral RNA has been mostly attributed to the p110 isoform. Our studies with IAV show that p110 isoform of ADAR1 is antiviral, as evidenced by increased viral titers in p110 KOs. It should be noted that the p110 KOs showed low levels of p110 induction upon treatment with IFN likely due to a cryptic promoter, as previously reported (Figure 2B) (Chung et al., 2018; Nachmani et al., 2014). Despite low levels of p110, replication of different IAV strains was 5-10 fold higher in p110 KOs to CTRL A549s (Figure 2C – Figure Supplement 2A). In agreement, prior studies suggest that p110 is relocalized from the nucleoplasm to the nucleolus during IAV infection, perhaps to prevent antiviral activity (de Chassey et al., 2013; Emmott et al., 2010). Future studies will determine if the RNA editing activity or other functional domains in p110 is critical for antiviral activity.

Recently, several studies have focused on understanding the role of ADAR1 in regulating cell intrinsic immunity (Lamers et al., 2019). Loss of ADAR1 or p150 results in embryonic lethality in mice, due to aberrant upregulation of ISGs and increased apoptosis in hematopoietic cells (Liddicoat et al., 2015; Pestal et al., 2015; Mannion et al., 2014; Ward et al., 2011). Embryonic lethality from ADAR1 deficiency can be rescued by concurrent deletion of RLR components MDA5 or MAVS, as doubly deficient mice show decreased ISG expression; however, ADAR1/MDA5 DKO mice display early lethality after birth (Pestal et al., 2015; Liddicoat et al., 2015; Mannion et al., 2014). In agreement, our studies in p150 KOs showed elevated basal IFN-β expression, which was reduced with concurrent deletion of MDA5, demonstrating the role of p150 in suppressing MDA5 mediated sensing of endogenous ligands under physiological conditions (Figure 3A and 3C). Despite displaying reduced basal IFN-β expression, the p150/MDA5 DKOs were unable to support robust IAV replication, suggesting that the role of p150 during IAV replication is independent of MDA5 suppression. Thus, our studies reveal an additional role for p150 in the suppression of RLR signaling during viral infection.

In addition to genetic deletion studies in mice, previous studies show increased ISG expression upon deletion of ADAR1 or p150 in transformed cell lines (Li et al., 2017; Chung et al., 2018; Li et al., 2012; George et al., 2016). Furthermore, treatment of these cells with exogenous type I IFN enhanced endogenous dsRNA induced antiviral responses, including activation of dsRNA protein kinase (PKR) and the OAS-RNase L pathway, resulting in shutdown of global protein translation by eIF2*α* phosphorylation and indiscriminate RNA degradation, respectively (Chung et al., 2018; Li et al., 2017; Pfaller et al., 2018). In this system, siRNA KD of PKR in ADAR1 KO cells or concurrent deletion of ADAR1 with RNase L resulted in restoration of protein translation and abrogation of RNA degradation, respectively. However, in our studies in p150 KOs and p150/MDA5 DKOs without addition of exogenous IFN, VSV replication was mostly unaffected, suggesting that restriction of IAV replication occurred via a mechanism independent of PKR and RNase L (Figure 2C and 3D). These results are in agreement with prior studies demonstrating that VSV replication is unaffected by loss of ADAR1 alone (Ward et al., 2011).

While several murine genetic studies have described the relationship between ADAR1 and MDA5, these studies were unable to fully examine the relationship between ADAR1 and RIG-I in mice due to the unrelated embryonic lethality associated with RIG-I ablation (Pestal et al., 2015). A study in human embryonic kidney cells (HEK293) showed that over expression of p110 or p150 resulted in decreased association of pI:C with RIG-I, suggesting that ADAR1 may prevent the sensing of dsRNA by RIG-I (Yang et al., 2014). Our studies in p150/MDA5/RIG-I or p150/MDA5/MAVS TKOs showed restoration of IAV replication and reduced IFN-β expression to levels similar to CTRL A549s, demonstrating that the p150 isoform is critical for suppressing RIG-I signaling during IAV infection (Figure 5D). We observed sustained IFN-β expression in p150/MDA5 DKOs, whereas IFN-β expression declined after initial induction in CTRL A549s (Figure 4C-D). Furthermore, differences in IAV replication between p150/MDA5 DKOs and CTRL A549s, were more pronounced at later times during infection (Figure Supplement 3B). Together, these results indicate that p150 acts as a negative regulator for ramping down RIG-I signaling during viral infection.

In addition to viral restriction through IRF3-mediated transcriptional upregulation of IFN-β and ISGs, viral spread can be restricted by cell death through cell intrinsic and extrinsic pathways (Elmore 2007). The RIPA pathway is a cell intrinsic apoptosis process that is triggered by activation of RLRs and subsequent linear polyubiqutinylation of IRF3 (Chattopadhyay et al., 2016). Polyubiquitinylated IRF3 in complex with Bax associates on the mitochondrial membrane and initiates apoptosis by promoting the release of Cytochrome C. Prior studies in ADAR1 deficient mice showed increased cell death in hematopoietic cell compartments along with elevated ISG expression (Qiu et al., 2013; Hartner et al., 2009). Moreover, measles virus infection of ADAR1 KD cells showed increased apoptosis (Toth et al., 2009). In our studies, ADAR1 KOs, p150 KOs, and p150/MDA5 DKOs showed elevated apoptosis upon IAV infection or stimulation with RLR agonist as compared to CTRL A549 (Figure 4F-H – Figure Supplement 4B-E). These results demonstrate that loss of p150 predisposes cells to RLR induced apoptosis. As further validation, concurrent deletion of Bax with p150 as well as p150/MDA5/RIG-I and p150/MDA5/MAVS TKOs showed decreased apoptosis upon vRNA transfection or IAV infection, demonstrating that p150 suppresses apoptosis via the RIPA pathway (Figure 6A and Figure 5C). A recent study shows that concurrent deletion of Bax with ADAR1 is not sufficient to rescue mice from embryonic lethality (Walkley et al., 2019). In our studies with p150/Bax DKOs, we observed reduced apoptosis yet high IFN-β expression upon RLR stimulation (Figure 6A-B). As such, treatment of p150/Bax DKOs with Ruxolitinib, a JAK inhibitor, resulted in significantly higher viral replication, demonstrating that restriction of IAV replication in p150 deficient cells occurred due to both elevated IFN receptor signaling and increased apoptosis via the RIPA pathway.

Both editing dependent and independent ADAR1 functions have been implicated in the suppression of innate immunity. Previous studies show that the RNA binding functions and editing functions of p110 and p150 were necessary for suppressing endogenous dsRNA or pI:C induced RLR signaling (Liddicoat et al., 2015; Yang et al., 2014). Similarly, in the context of measles virus infection, ADAR1 KO cells complemented with a catalytic mutant of p150 showed a modest increase in viral replication as compared to control ADAR1 KO, whereas wildtype p150 completely restored viral replication (Pfaller et al., 2018). Moreover, murine genetic studies also highlight the importance of ADAR1 mediated editing of endogenous dsRNA in the suppression of MDA5 activation. However, in another study, overexpression of p110 or p150 catalytic mutants was sufficient to suppress RIG-I activation upon SeV infection (Yang et al., 2014). Our studies with p150/MDA5 DKOs complemented with different p150 mutants demonstrated that the RNA binding ability of p150 but not the catalytic activity was critical for suppression of RIG-I signaling. We hypothesize that p150 may function upstream of RIG-I activation, possibly by binding and sequestering cellular or viral RNA to prevent sustained RIG-I sensing and activation.

Our data shows that the role of p150 in promoting viral replication is specific to IAV, as VSV replication was unaffected in p150 deficient cells (Fig 3D). The NS1 of IAV has been show to interact with ADAR1 in yeast-2-hybrid screening and immunoprecipitation assays, as well as to increase the editing activity of ADAR1 in reporter assays (de Chassey et al., 2013; Ngamurulert et al., 2009). However, the significance of these interactions has yet to be determined. Importantly, NS1 has been shown to antagonize several host antiviral pathways, including activation of the RIG-I pathway via its interactions with the E3 ligases TRIM25 and RIPLET (Gack et al., 2009; Mibayashi et al., 2007; Rajsbaum et al., 2012). Despite the presence of this viral antagonist, we observed decreased IAV replication in p150 deficient cells due to increased RIG-I signaling, suggesting that NS1 is likely incapable of suppressing the amplification of RIG-I signaling in cells lacking p150 (Fig 5B). Future studies will determine the importance of NS1:ADAR1 interactions in suppression of the RIG-I pathway.

In conclusion, our studies highlight the importance of p150 as a negative regulator of RLR signaling. p150 prevents endogenous dsRNA mediated activation of the innate immune sensor MDA5 under basal conditions (Liddicoat et al., 2015; Pestal et al., 2015; Chung et al., 2018). Our studies demonstrate that p150 also functions to dampen host antiviral signaling via suppression of RIG-I during IAV infection. This dampening of RIG-I mediated signaling by p150 enables efficient IAV replication. Future studies are required to determine the exact mechanism of p150 mediated suppression of the RIG-I pathway and the significance of NS1-p150 interactions in this process.

## Materials and Methods

### Cell Culture and Viruses

Human lung epithelial (A549) and human embryonic kidney (HEK293) cells were cultured in DMEM supplemented with 10% fetal bovine serum (FBS) and 1% penicillin/streptomycin (P/S). Madin-Darby canine kidney (MDCK) cells were cultured in MEM supplemented with 10% FBS and 1% P/S. Dr. Adolfo Garcia-Sastre at the Icahn School of Medicine at Mount Sinai, NY kindly provided the IAV strains A/Puerto Rico/8/1934 (H1N1), low pathogenic version of A/Vietnam/1203/2004 (H5N1), and A/Hong Kong/1/1968 (H3N2). IAV strains were grown in 10-day old specific pathogen-free eggs (Charles River) and titered by standard plaque assay on MDCK cells, using 2.4% Avicel RC-581 (a gift from FMC BioPolmer, Philadelphia, PA). Two days post infection plaques were quantified via crystal violet staining. Vesicular stomatitis virus expressing GFP (VSV) was kindly provided by Dr. Glenn Barber at the University of Miami, FL (Stojdl et al., 2003). VSV viruses were grown in Vero cells and titered by plaque assay with 1% methylcellulose (Sigma).

### Generation of CRISPR KO and cDNA Complemented Cells

ADAR1 KO A549s were generated using the lentiCRISPR v2 single vector system, which has been previously described (Sanjana et al., 2014). Oligonucleotides were annealed and cloned into the lentiCRISPR v2 vector through BsmBI sites (Oligonucleotide table). A549s were transduced with lentivirus expressing specific sgRNA and selected with 2 μg/ml puromycin for 14 days. CTRL A549s were generated through the same method using the lentiCRISPR v2 vector without sgRNA. To generate clonal KO A549s, puromycin selected population were seeded at ∼100-150 cells per 15cm plates and individual colonies were isolated using cloning cylinders. Subsequently, gDNA was extracted from isolated KO cells using the Blood and Cell Culture Mini kit (Qiagin) per the manufacturers specification and the targeted region was PCR amplified using EconoTaq and gene specific primers. PCR products were cloned into pGEM-T vector and approximated 12-15 individual bacterial colonies were sequenced (University of Chicago DNA sequencing facility). Sequences were analyzed by SeqMan program (LaserGene).

To generate isoform specific KO A549s, WT A549s were transduced with lentiCRISPR v2 containing sgRNA targeting a region 200-300 nucleotides upstream of transcription start site and lentiCRISPR v2-GFP containing a sgRNA downstream of the transcription start site. Cells were selected with 2 μg/ml puromycin for 14 days and then selected for GFP expression by single-cell sorting, to ensure the presence of both sgRNA. Individual KO clones were identified by PCR analysis of targeted region and confirmed by western blot for ADAR1 expression.

To generate p150/MDA5 DKOs, clonal p150 KOs were transduced with a lentiCRISPR v2-hygro vector containing gene specific sgRNA followed by selection with 600 μg/ml hygromycin for 14 days. For the p150/Bax DKOs, clonal p150 KOs were transduced with a lentiCRISPR v2-neo vector containing the gene specific sgRNA followed by selection in 600 μg/ml neomycin for 14 days. The RIG-I/MDA5/p150 and MAVS/MDA5/p150 TKO A549s were generated by transducing the p150/MDA5 DKOs with the lentiCRISPR v2-neo vector containing gene specific sgRNA followed by selection in 600 μg/ml neomycin for 14 days. Individual KO clones were isolated as described above and screened by western blot and Sanger sequencing.

To generate complemented cell lines, p150 cDNA was cloned into the pLX304 lentivirus vector. p150 KO and p150/MDA5 DKOs were transduced with lentivirus carrying either empty vector or p150 cDNA, and selected with 15 μg/ml blasticidin for 14 days. Polyclonal population were seeded at limiting dilutions and individual clones were isolated and confirmed via western. After selection, complemented clones were maintained in 5 μg/ml blasticidin.

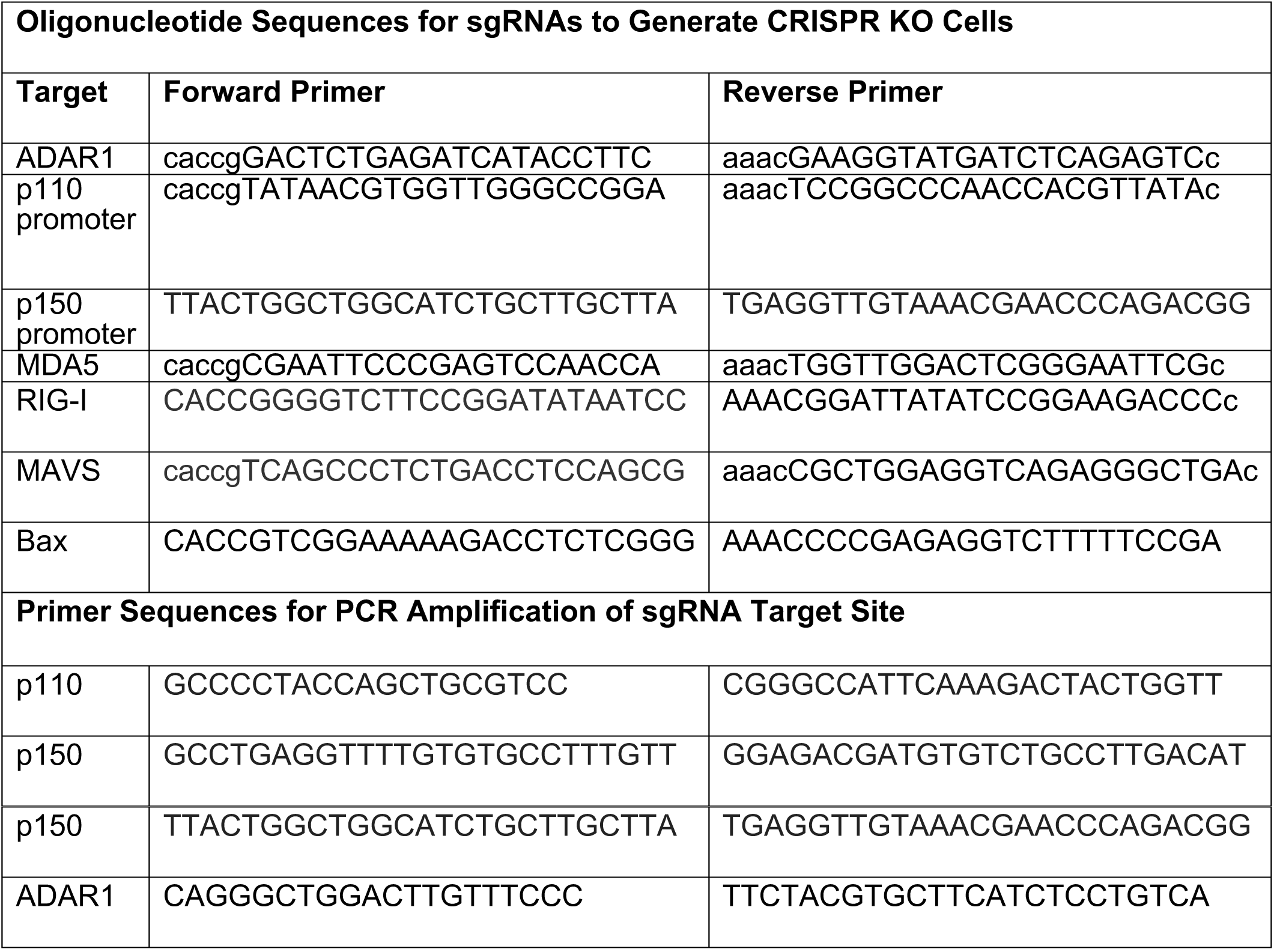

### Virus Infections

To assess viral replication, CTRL A549s or KOs were seeded at a density of 1.2×10^6^ cells per well in triplicate in a 6-well dish. For IAV infection, cells were washed twice with phosphate buffer saline (PBS) and 500 μl of infection media (DMEM supplemented with 0.2% bovine serum albumin (BSA) and 1μg/ml TPCK-treated trypsin (T1426, Sigma)) was added per well. Cells were infected at indicated MOI based on cell numbers determined prior to infection and incubated for 1 hour at 37°C. After the 1 hour incubation, the inoculum was removed and the cells were washed twice in PBS and placed in 2 ml of fresh infection media. Supernatants were harvested at the indicated times and titered by plaque assay in MDCK cells (Han et al., 2018). For VSV, infections were performed as described above with DMEM supplemented with 2% FBS. VSV titers were assessed by plaque assay in Vero cells using 1% methylcellulose. To examine viral replication following inhibition of IFN signaling, cells were seeded at a density of 1.2×10^6^ cells per well in a 6-well dish in DMEM supplemented with 10% FBS, 1% P/S, and either DMSO or 0.2 μM Ruxolitinib (INCB018424 Selleckchem). IAV infection was performed as described above using infection media containing DMSO or 0.2 μM Ruxolitinib. Viral titers were assessed as described above. All infection experiments were performed with biological triplicates and data is presented as the mean titer of the triplicate samples(±SD).

For western blotting and qPCR, cells were seeded at a density of 1.2×10^6^ cells per well in a 6-well dish. Cell numbers were calculated the next day prior to infection. Cells were mock treated or inoculated with H1N1 at MOI of 1 or SeV at a 1:100 dilution in DMEM supplemented with 0.2% BSA and 1% P/S at 37°C. Cells were harvested at the indicated time points for RNA or protein isolation.

### QPCR Analysis of Host Gene Expression

Total RNA from pooled triplicates was isolated by TRIzol (Invitrogen) extraction method. Residual genomic DNA contamination was removed by treatment with recombinant DNAse I (Roche). cDNA was generated using Superscript II reverse transcriptase and Oligo d(T) (Invitrogen). Using SYBR Green PCR Master Mix (Applied Biosystems), qPCR was performed with technical duplicates and gene specific primers. Tubulin was used as an endogenous house housekeeping gene to calculate delta delta cycle thresholds. Results are represented as fold expression relative to mock vector CTRL A549s.

RT and qPCR primers: hIFN-β: forward TCTGGCACAACAGGTAGTAGGC and reverse GAGAAGCACAACAGGAGAGCAA; hTub: forward GCCTGGACCACAAGTTTGAC and reverse TGAAATTCTGGGAGCATGAC.

### Western Blot Analysis

To confirm ADAR1 isoform specific KO and MDA5 KOs, CTRL A549s and various KOs cells were treated with 1000U/ml of universal type I IFN (PBL Assay Science) for 24 hours and lysed for western blot analysis. To confirm RIG-I KO, vector control and KO cells were infected with SeV for 16 hours and then lysed for western blot analysis. Whole cell lysates for western blot analysis was prepared using RIPA buffer (50 mM Tris pH 7.8, 150 mM NaCl, 0.1% SDS, 0.5% Sodium deoxycholate, 1% Triton X-100) containing protease inhibitors and quantified by Bradford protein assay. Approximately 50-80 μg of total protein was loaded on to SDS gel. Antibodies used for western blot analysis: ADAR1 (Abcam 88574; Santa Cruz sc-271854), MDA5 (Alexis Biochemicals AT113), RIG-I ((1C3 clone) (Nistal-Villan et al., 2010)), IRF3 (Santa Cruz sc-9082), V5 (Bio-Rad), cleaved PARP (Cell Signaling 5625), total PARP (Cell Signaling 9542), and Ku (#K2882 Sigma).

### RLR agonist transfection

For vRNA transfection, IAV vRNA from H1N1 was extracted using the QIAamp Viral RNA Mini Kit per the manufacturer’s recommendations. Cells were seeded at a density of 1.2×10^6^ cells per well in a 6-well dish a day prior to transfection. Cells were transfected with vRNA using polyethylenimine reagent (PEI) in OptiMem (OMEM) and were collected at the indicated time points for RNA or protein extraction. For low molecular weight (LMW) and high molecular weight (HMW) poly(I:C) (pI:C) transfections, 1μg of LMW or HMW pI:C (Invivogen) was transfected into cells

For siRNA transfection, cells were seeded at a density of 1.5×10^5^ cells per well in a 12-well dish. 50 nM pooled siRNA (Dharmacon) was transfected using RNAiMAx reagent (2.4 μl per well). At 48 hours post siRNA transfection, cells were transfected with H1N1 vRNA and cell lysates were collected 24 hours post vRNA transfection for western blot.

### IFN-β reporter assay

p150 KO 293s was transfected with a plasmid mixture containing p150 (100ng), IFN-β-Firefly reporter (100ng), and SV40 Renilla (50ng) using PEI (1:5 DNA:PEI ratio). A GFP expressing plasmid was included in place of p150 as controls. IFN-β-Firefly reporter activity was induced by infecting cells with SeV (1:100) at 6 hours post transfection. At 48 hours post transfection, cells were lysed in 1x Passive Lysis Buffer (Promega), and firefly and renilla luciferase activity in the lysates were measured using Dual-Luciferase Reporter Assay System (Promega) in GloMax 20/20 luminometer (Promega). Data from biological triplicates is presented as percent luciferase activity relative to GFP + SeV control.

### ANNEXIN V Staining

ADAR1 KOs were seeded at a density of 1.0×10^6^ cells per well in a 6-well dish. The following day, cells were transfected with H1N1 vRNA as described above. At 40 hours post transfection, cells were stained with Annexin V-PE (BD Pharmingen) per the manufacturer’s recommendations. Annexin V staining was analyzed by flow cytometer. Data analysis was performed using FlowJo Software.

## Acknowledgements

We would like to thank Dr. Adolfo Garcia-Sastre (Icahn School of Medicine) for sharing numerous reagents. Olivia Vogel and Julianna Han were partly supported by the NIH Molecular and Cellular Biology training program at The University of Chicago (T32GM007183); Julianna Han was partly supported the NIH Diversity Supplement (R01AI123359-02S1). Dr. Balaji Manicassamy is supported by NIAID grants (R01AI123359 and R01AI127775).

## Author Contributions

OV, JTP and BM conceived and designed the study. OV, JTP and BM generated the data shown in all figures. JH helped with the generation of KO cells and expression plasmids. CL performed the flow cytometry analysis for Annexin V staining. SM provided scientific input on the project. OV, JTP and BM wrote the manuscript. All authors approved the manuscript.

---

**Figure Supplement 1.**
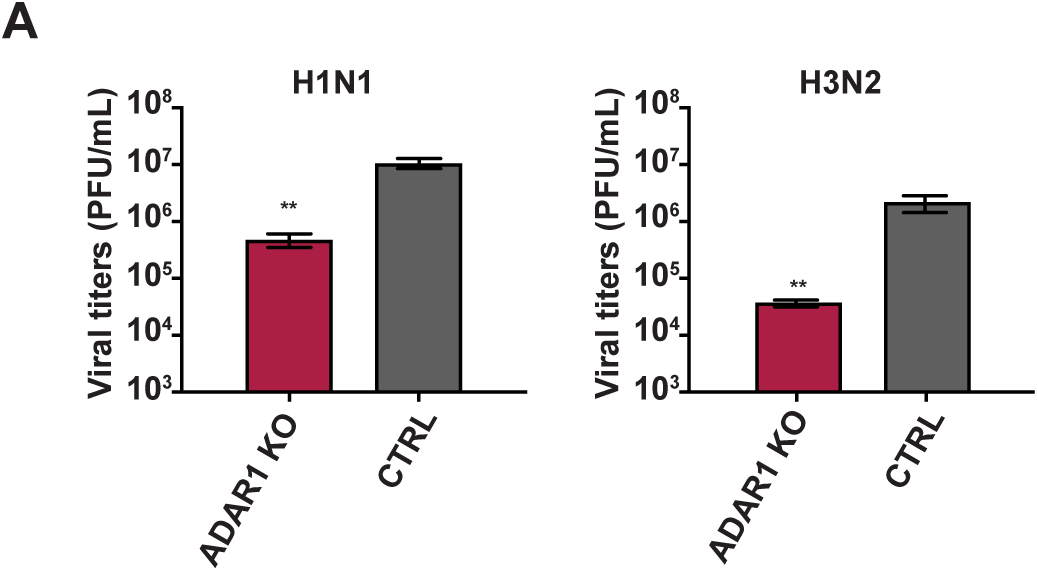
IAV strains replicate poorly in ADAR1 Kos. (A) ADAR1 KOs and CTRL A549s were infected with H1N1 (MOI = 0.01) and H3N2 (MOI = 0.01). Viral titers were measured at 48 hours. Data are represented as mean titer of triplicate samples ± SD. * denotes p-value *≤* 0.5. ** denotes p-value *≤* 0.01. *** denotes p-value *≤* 0.001. NS denotes p-value *≥* 0.05. Data are representative of at least three independent experiments.

**Figure Supplement 2.**
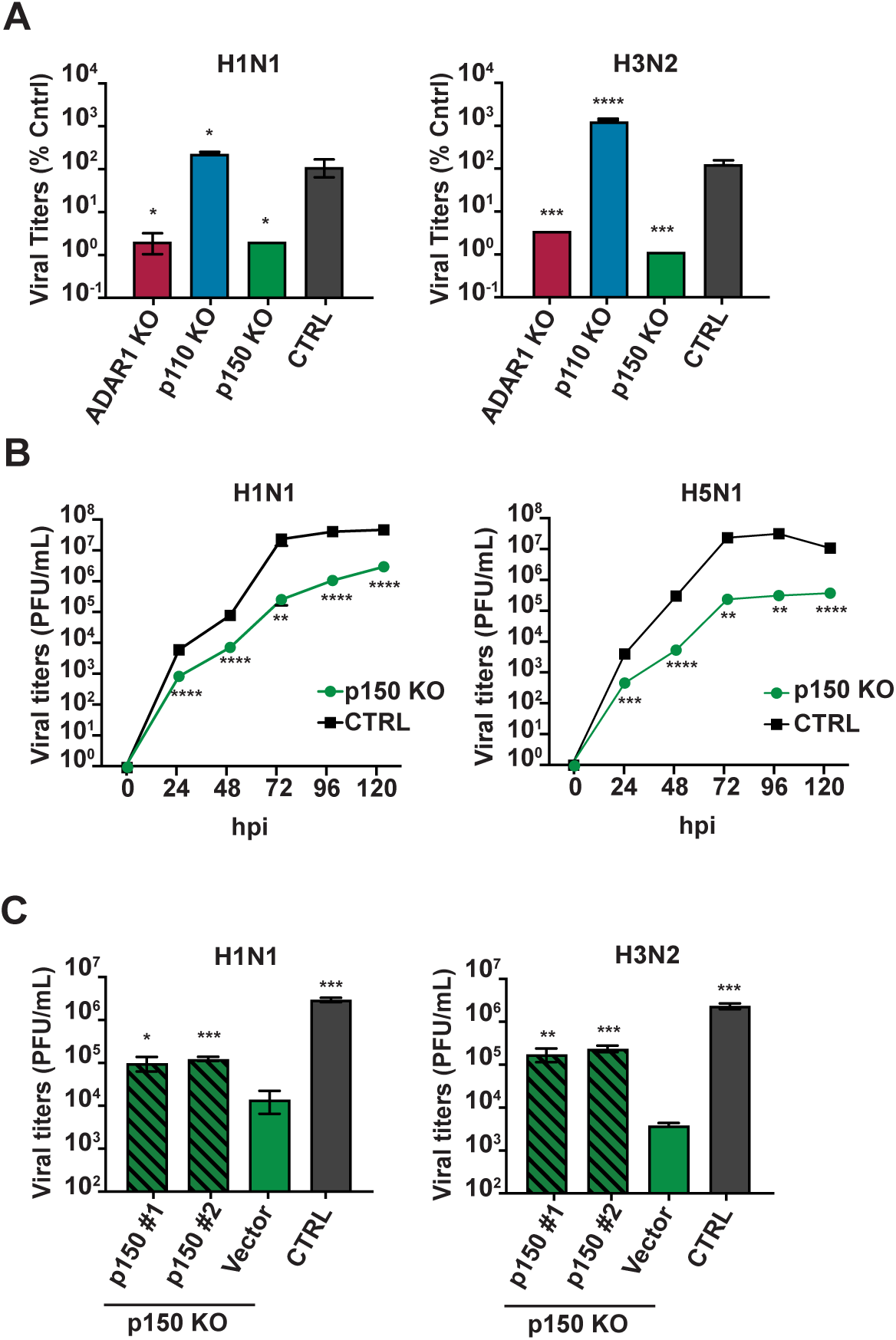
p150 isoform of ADAR1 is critical for IAV replication. (A) Assessment of viral replication in various KO cells lacking ADAR1 isoforms. ADAR1 KOs, isoform specific KOs, and CTRL A549s were infected with H1N1 (MOI = 0.01) and H3N2 (MOI = 0.01) and viral titers were measured at 48 hpi. (B) p150 KOs and CTRL A549s were infected with H1N1 (MOI = 0.01) and H5N1 (MOI = 0.001) and viral titers were measured at the indicated times post infection. (C) Two clones of p150 KOs complemented with wildtype p150, empty vector p150 KOs, and CTRL A549s were infected with H1N1 (MOI = 0.01) and H3N2 (MOI = 0.01) and viral titers were measured at 48 hpi. Data are represented as mean titer of triplicate samples ± SD. * denotes p-value *≤* 0.5. ** denotes p-value *≤* 0.01. *** denotes p-value *≤* 0.001. NS denotes p-value *≥* 0.05. Data are representative of at least three independent experiments.

**Figure Supplement 3.**
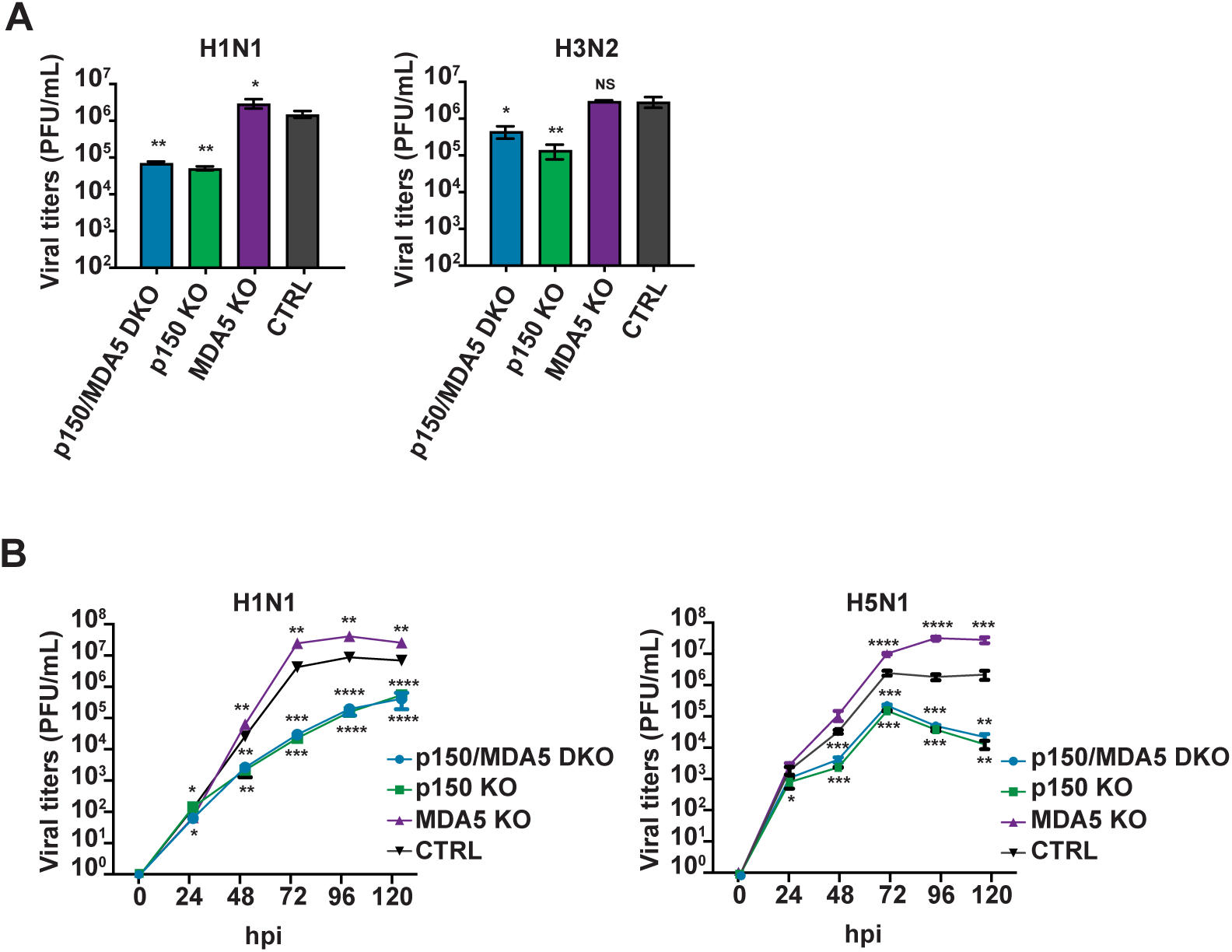
p150 enhances IAV replication independent of its ability to suppress MDA5. (A) Assessment of viral replication in various KOs. p150/MDA5 DKOs, p150 KOs, MDA5 KOs, and CTRL A549s were infected with H1N1 (MOI = 0.01) and H3N2 (MOI = 0.01). Viral titers were measured at 48 hours. (B) Viral replication kinetics in various KOs. p150/MDA5 DKOs, p150 KOs, MDA5 KOs, and CTRL A549s were infected with H1N1 (MOI = 0.01) and H3N2 (MOI = 0.01). Viral titers were measured at the indicated time points post infection. Data are represented as mean titer of triplicate samples ± SD. * denotes p-value *≤* 0.5. ** denotes p-value *≤* 0.01. *** denotes p-value *≤* 0.001. NS denotes p-value *≥* 0.05. Data are representative of at least three independent experiments.

**Figure Supplement 4.**
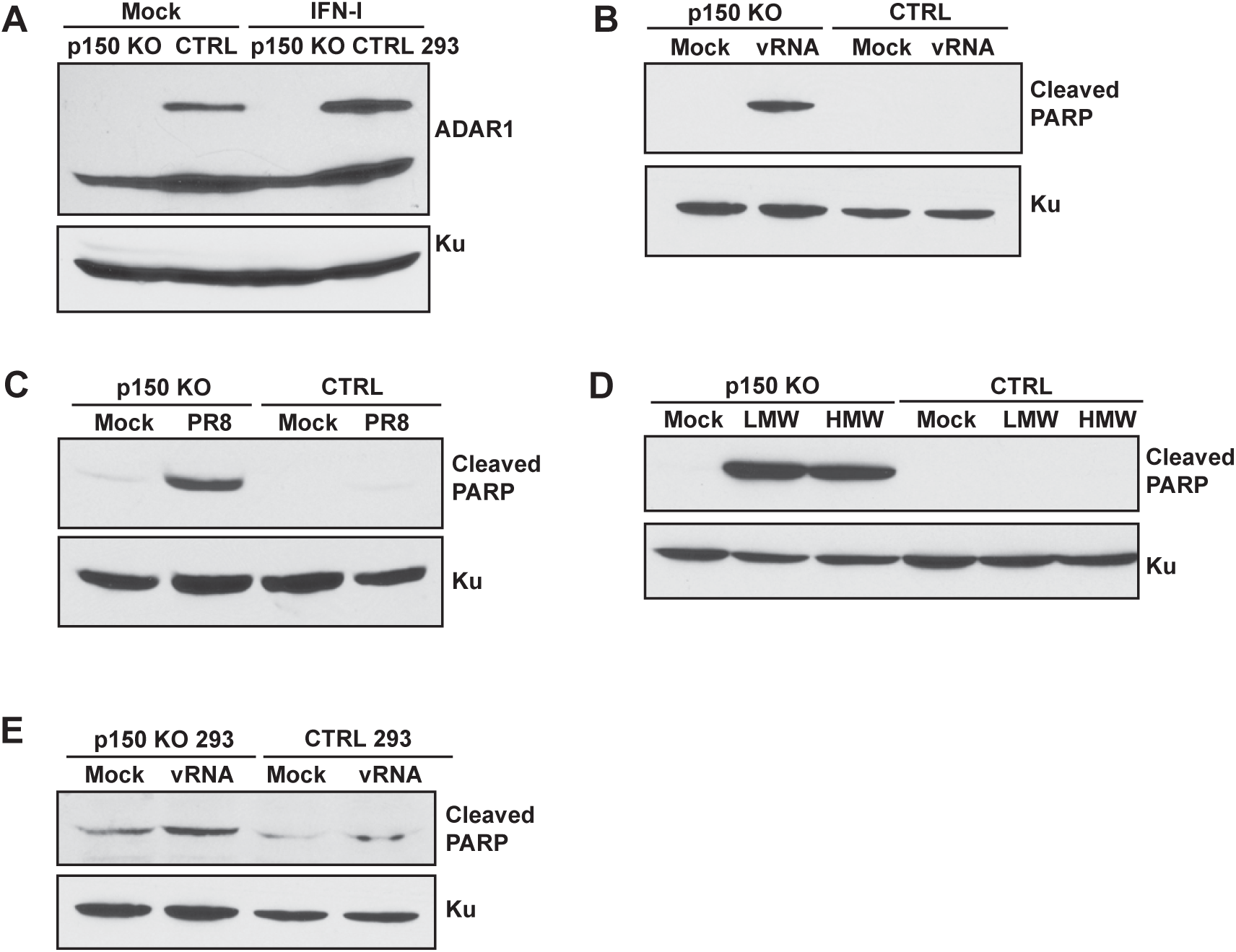
p150 KOs show increased PARP cleavage following stimulation of RLRs. (A) Western blot analysis of ADAR1 expression in p150 KOs and CTRL 293s. p150 KOs, and CTRL 293s were mock treated or treated with IFN for 24 hours and expression of ADAR1 was examined by western blot. Expression of Ku is shown as a loading control. (B-D) Western blot analysis of PARP cleavage upon RLR stimulation. (B) PARP cleavage in p150 KOs following IAV vRNA transfection. p150 KOs and CTRL A549s were transfected with H1N1 vRNA. Lysates were collected at 24 hours post transfection. (C) PARP cleavage p150 KOs following H1N1 infection. p150 KOs and CTRL A549s were infected with H1N1 (MOI =1). Lysates were collected at 40 hours post infection. (D) PARP cleavage in p150 KO following poly I:C transfection. p150 KOs and CTRL A549s were transfected with LMW or HMW pI:C. Lysates were collected at 24 hours post transfection. (E) PARP cleavage in p150 KO 293s following IAV vRNA transfection. p150 KOs and CTRL 293s were transfected with H1N1 vRNA. Lysates were collected at 24 hours post transfection.

